# *Scn2a*-deficient mice exhibit serotonergic dysfunction and altered behavioral responses to psilocybin

**DOI:** 10.64898/2026.11.03.764054

**Authors:** Brody A. Deming, Jingliang Zhang, Josephine Brown-Leung, Manasi Halurkar, Akila D. Abeyaratna, Rineet Ranga, Keeley Farmer, Deepti Ramasamy, Jason Cannon, Yang Yang

**Author notes:** **To whom correspondence should be addressed:** Yang Yang, PhD; Borch Department of Medicinal Chemistry and Molecular Pharmacology, College of Pharmacy, Purdue University, 207 S Martin Jischke Dr, West Lafayette, IN 47907.

## Abstract

Autism spectrum disorder (ASD) is a prominent neurodevelopmental disorder affecting approximately 3.2% of children in the United States. Although ASD has traditionally been considered a polygenic disorder, advances in genetic testing have identified individual genes that can contribute to its development. Large-scale whole-exome sequencing has identified variants in *SCN2A*, which encodes the alpha subunit of the voltage-gated sodium channel Nav1.2, as a leading monogenic cause of ASD. Social deficits are a core feature of ASD, and the serotonergic system has been implicated in the regulation of social behavior. Recent studies have demonstrated that serotonergic psychedelics, such as psilocybin, can increase sociability in mouse models of ASD. However, whether underlying genetic alterations associated with ASD influence behavioral responses to psilocybin remains poorly understood. Therefore, we investigated how *Scn2a* deficiency affects behavioral responses to psilocybin and whether these differences are accompanied by alterations in the serotonergic system. We found that *Scn2a*-deficient mice exhibited differential responses to psilocybin compared to wild-type (WT) mice. Specifically, psilocybin increased sociability in *Scn2a*-deficient mice without affecting anxiety-like behavior, whereas WT mice exhibited decreased anxiety-like behavior without increased sociability. Further behavioral evaluation revealed a reduced head twitch response in *Scn2a*-deficient mice following psilocybin administration compared to WT mice. To further investigate these differences, we examined serotonin levels and receptor expression in the medial prefrontal cortex (mPFC), a brain region implicated in social behavior and psychedelic action. Although baseline serotonin levels in the mPFC did not differ between genotypes, *Scn2a*-deficient mice exhibited altered serotonin receptor expression. We further characterized the serotonergic system by evaluating neuronal activity in the Dorsal Raphe Nucleus, a major source of serotonergic projections to the mPFC, and found increased spontaneous firing in *Scn2a*-deficient mice. Together, these findings highlight serotonergic alterations associated with *SCN2A* deficiency and the potential for serotonergic interventions to alleviate social deficits associated with *SCN2A*-related ASD.

## INTRODUCTION

Psychedelics are a class of compounds that profoundly alter an individual’s perception of the world [1, 2]. Research surrounding these molecules has suggested their potential for treating a range of disorders, including autism spectrum disorder (ASD) [3–5]. The prevalence of ASD has increased in recent years, with approximately 3.2% of children in the United States identified as having ASD [6]. Large-scale whole-exome sequencing has identified variants in the gene *SCN2A*, which encodes the voltage-gated sodium channel Nav1.2, as one of the leading monogenic causes of ASD [7, 8]. ASD presents with a range of symptoms, with alterations in social behavior being one of its core features [9]. Preclinical models that reliably recapitulate these social deficits are important for understanding *SCN2A*-related ASD and evaluating potential therapeutics.

Although *Scn2a^+/-^* mice have been used to model *SCN2A*-related ASD, they do not consistently exhibit the pronounced social deficits associated with this disorder, which can limit their utility for studying social behavior and evaluating potential therapeutics [10–12]. To overcome this limitation, our lab previously developed a *Scn2a-*deficient mouse model (*Scn2a^gt/gt^*) by inserting a gene-trap cassette into the *Scn2a* gene, resulting in approximately 25-30% expression of Nav1.2 and robust ASD-relevant behavioral phenotypes [13–16]. The pronounced social deficits observed in these mice provide an opportunity to evaluate potential treatments targeting social behavior. Early work evaluating the therapeutic potential of psychedelics suggested that they could enhance social behavior in individuals with ASD [4, 17–19]. More recent work has extended these findings, demonstrating that psychedelics modulate social cognition in humans and can influence social behavior in preclinical models [20–24].

Psilocybin (Psi), a naturally occurring psychedelic compound found in “magic mushrooms”, is converted to its active metabolite, psilocin, which exerts its psychedelic effects primarily through activation of the 5-HT2A receptor [25, 26]. However, psilocin also interacts with several other serotonin (5-HT) receptor subtypes, many of which have been implicated in the regulation of social behavior [27–35]. Despite the importance of serotonergic signaling in social behavior, whether the serotonergic system is altered in *SCN2A*-related ASD remains unknown. Furthermore, whether *Scn2a* deficiency influences behavioral responses to Psi has yet to be determined.

In this study, we investigated the genotype-dependent behavioral effects of Psi in WT and *Scn2a^gt/gt^* mice and whether alterations in the serotonergic system associated with *Scn2a* deficiency could help explain these differences. We found that Psi acutely increased sociability in *Scn2a^gt/gt^* mice, an effect not observed in wild-type (WT) mice. This increase in sociability was not accompanied by significant changes in locomotor activity. In contrast, Psi increased time spent in the center of the open field in WT mice but not *Scn2a^gt/gt^* mice. Additionally, *Scn2a^gt/gt^* mice exhibited a reduced head twitch response (HTR) following Psi administration, suggesting genotype-dependent differences in behavioral responses to Psi. Given these differences, we next evaluated whether the serotonergic system was altered in *Scn2a^gt/gt^* mice. qPCR and ultra-high-performance liquid chromatography with electrochemical detection (UHPLC-ECD) revealed minimal changes in 5-HT receptor mRNA expression and no differences in 5-HT, 5-hydroxyindoleacetic acid (5-HIAA), or 5-HT turnover in the medial prefrontal cortex (mPFC). However, Western blot analyses revealed altered expression of multiple serotonin receptors and the serotonin transporter (SERT) in *Scn2a^gt/gt^* mice. Given that the Dorsal Raphe Nucleus (DRN) is a major source of serotonergic projections to the mPFC, we next evaluated spontaneous neuronal activity in this region using patch-clamp electrophysiology and found increased spontaneous firing in *Scn2a^gt/gt^* mice, suggesting that *Scn2a* deficiency may alter serotonergic circuit activity. Together, these findings highlight the complexity of serotonergic signaling and the potential of targeting the serotonergic system as a therapeutic approach for reducing social abnormalities associated with *SCN2A*-related ASD.

## RESULTS

### Psilocybin acutely increases sociability in *Scn2a*-deficient mice but not in WT mice

Given the important role 5-HT plays in modulating social behavior and the rapidly expanding field of psychedelic medicine [3, 36], we aimed to determine whether Psi could increase social behavior in *Scn2a^gt/gt^* mice. *Scn2a^gt/gt^* and WT mice underwent a three-chamber assay to assess social approach toward a novel mouse and a stranger interaction assay to evaluate direct social interactions in a more naturalistic setting. Mice were first injected with saline (Veh) to establish baseline social behavior and then injected with Psi one week later. Mice were tested again one week after Psi administration to determine whether the effects of Psi persisted (**Figure 1A**). Consistent with our previous observations [14, 37], at baseline, *Scn2a^gt/gt^* mice exhibited severe social deficits, spending less time in the social zone and having fewer interactions with the novel mouse compared to WT mice (Time in social zone: WT: 126.4 ± 11.16 s; *Scn2a^gt/gt^*: 48.30 ± 18.74 s) (Number of interactions: WT: 14.30 ± 1.85, *Scn2a^gt/gt^*: 1.80 ± 1.09). Thirty minutes after Psi administration, we observed that *Scn2a^gt/gt^* mice had an increase in the time spent in the social zone (*Scn2a^gt/gt^*: 133.1 ± 33.67 s) and the number of interactions with the novel mouse (*Scn2a^gt/gt^*: 14.30 ± 4.60). Interestingly, the prosocial effect of Psi was not observed in WT mice, consistent with previous findings in WT mice [38, 39]. One week after Psi injection, *Scn2a^gt/gt^* mice returned to baseline behavior (Time in social zone: *Scn2a^gt/gt^*: 36.02 ± 15.44 s, # of interactions: *Scn2a^gt/gt^*: 2.10 ± 1.39), suggesting that the prosocial effect of Psi is acute (**Figures 1B**, **1C_i_, and 1C_ii_**).

**Figure 1:**
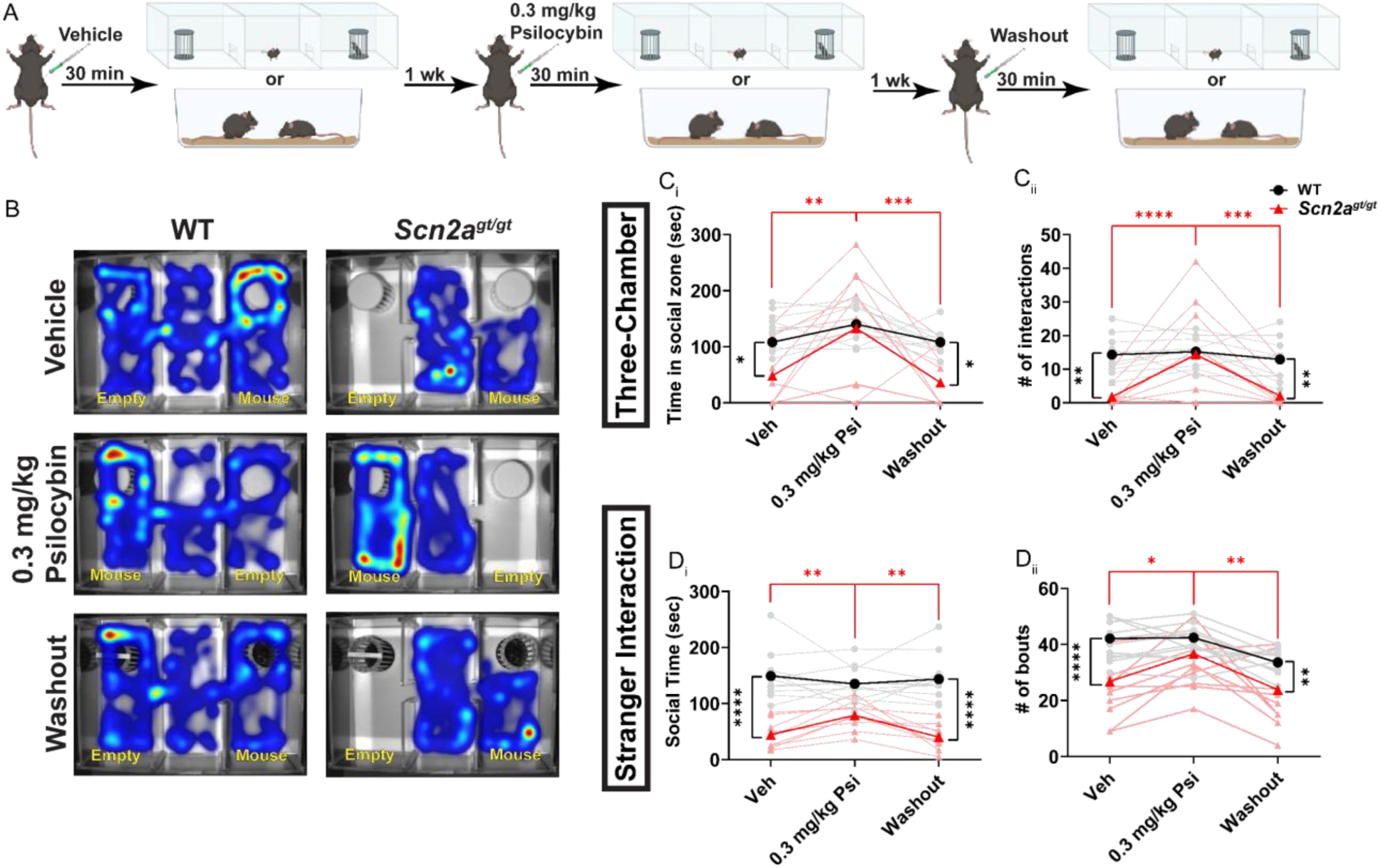
Psilocybin acutely increases sociability in *Scn2a*-deficient mice but not in WT mice. (A) Schematic of experimental design. (B) Representative heatmaps of WT and *Scn2a^gt/gt^* mice during a 5-minute three-chamber assay. (C_i_) Time WT and *Scn2a^gt/gt^* mice spent in the social zone during a 5-minute three-chamber assay 30 minutes after a vehicle or psilocybin injection. n= WT: 10, *Scn2a^gt/gt^*: 10. Two-way repeated-measures ANOVA with Bonferroni multiple comparisons: *, p<0.05. **, p<0.01, ***,p<0.001. (C_ii_) Number of interactions WT and *Scn2a^gt/gt^* mice had with a novel mouse during a 5-minute three-chamber assay 30 minutes after a vehicle or psilocybin injection. n= WT: 10, *Scn2a^gt/gt^*: 10. Two-way repeated-measures ANOVA with Bonferroni multiple comparisons: **, p<0.01, ***, p<0.001 ****, p<0.0001. (D_i_) Time WT and *Scn2a^gt/gt^* mice spent interacting with a novel mouse during the stranger interaction assay 30 minutes after a vehicle or psilocybin injection. n= WT: 11, *Scn2a^gt/gt^*: 9. Two-way repeated-measures ANOVA with Bonferroni multiple comparisons: **, p<0.01, ****,p<0.0001. (D_ii_) Number of bouts WT and *Scn2a^gt/gt^* mice had with a novel mouse during the stranger interaction assay 30 minutes after a vehicle or psilocybin injection. n= WT: 11, *Scn2a^gt/gt^*: 9. Two-way repeated-measures ANOVA with Bonferroni multiple comparisons: *, p<0.05. **, p<0.01, ****,p<0.0001.

We next examined whether these prosocial effects were also observed in a stranger interaction assay (**Figure 1A**). Consistent with our three-chamber assay, *Scn2a^gt/gt^* mice spent less time interacting with a novel mouse compared to WT mice (WT: 149.1 ± 12.99 s, *Scn2a^gt/gt^*: 44.33 ± 8.41 s) and also presented with a lower number of bouts with the novel mouse at baseline (WT: 40.55 ± 2.06, *Scn2a^gt/gt^*: 22.11 ± 3.34). Thirty minutes after Psi injection, we observed that *Scn2a^gt/gt^* mice had an increase in social time (*Scn2a^gt/gt^*: 79.00 ± 8.56 s) and number of bouts with the novel mouse (*Scn2a^gt/gt^*: 32.78 ± 3.36). These prosocial effects were not observed one week after Psi administration (Social time: *Scn2a^gt/gt^*: 40.22 ± 7.81 s, # of bouts *Scn2a^gt/gt^*: 21.11 ± 3.98) (**Figure 1D_i_ and 1D_ii_**). Together, these findings demonstrate that Psi acutely increases social behavior in *Scn2a^gt/gt^* mice, but these effects do not persist one week after treatment.

### Psilocybin decreases anxiety-like behavior in WT mice but not in *Scn2a*-deficient mice

Following the observed increase in sociability, we next examined whether Psi altered locomotor activity or anxiety-like behavior. WT and *Scn2a^gt/gt^* mice underwent an open field assay following Veh or Psi administration (**Figure 2A**). We found that Psi did not significantly alter the total distance traveled (WT: Veh: 2659 ± 131.5 cm, Psi: 3517 ± 208.4 cm, *Scn2a^gt/gt^*: Veh: 4167 ± 707.9 cm, Psi: 7017 ± 1762 cm) or average velocity (WT: Veh: 4.43 ± 0.22 cm/s, Psi: 5.86 ± 0.35 cm/s, *Scn2a^gt/gt^*: Veh: 6.96 ± 1.18 cm/s, Psi: 11.70 ± 2.94 cm/s) in WT or *Scn2a^gt/gt^* mice (**Figures 2C_i_ and C_ii_**), suggesting that the increased sociability observed in *Scn2a^gt/gt^* mice was not due to an overall increase in locomotor activity. Interestingly, WT mice injected with Psi showed an increase in time spent exploring the center of the open field arena (WT: Veh: 58.13 ± 12.75 s, Psi: 132.8 ± 20.64 s), while *Scn2a^gt/gt^* mice did not (*Scn2a^gt/gt^*: Veh: 34.26 ± 7.71 s, Psi: 44.06 ± 10.07 s) (**Figure 2D_i_**). Together, these findings suggest that the prosocial effect of Psi in *Scn2a^gt/gt^* mice is not driven by changes in locomotor activity and highlight distinct effects of Psi on sociability and anxiety-like behavior between *Scn2a^gt/gt^* and WT mice.

**Figure 2:**
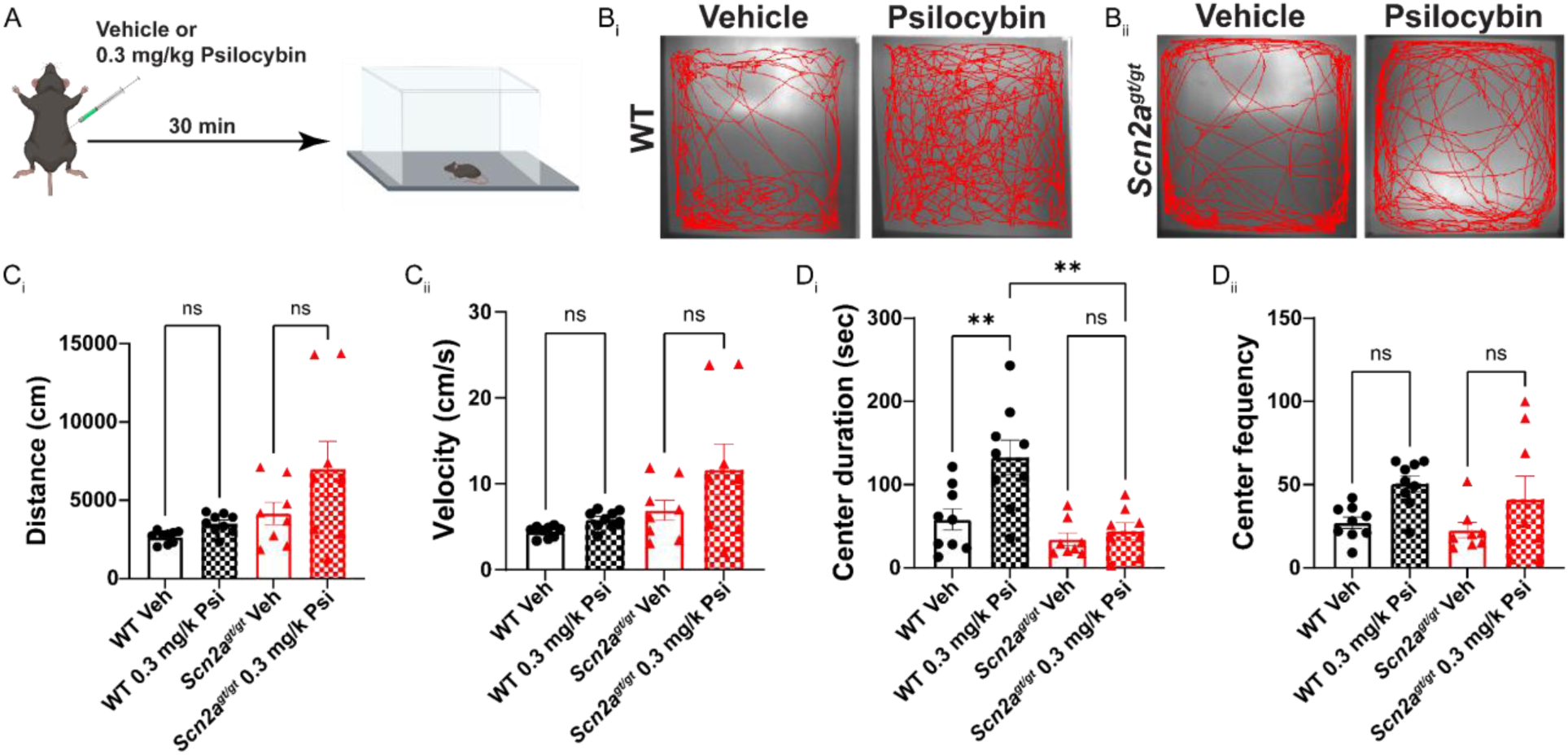
Psilocybin decreases anxiety-like behavior in WT mice but not in *Scn2a*-deficient mice. (A) Schematic of experimental design. (B_i_) Representative traces of the WT mice injected with vehicle or psilocybin during a 10 min open field assay. (B_ii_) Representative traces of the *Scn2a^gt/gt^* mice injected with vehicle or psilocybin during a 10 min open field assay. (C_i_) Distance traveled by WT or *Scn2a^gt/gt^* mice injected with vehicle or psilocybin during a 10 min open field assay. n= WT Veh: 9, WT Psi: 9, *Scn2a^gt/gt^* Veh: 8, *Scn2a^gt/gt^* Psi: 8. Two-way ANOVA with Bonferroni multiple comparisons: ns, p>0.05. (C_ii_) Average velocity of WT or *Scn2a^gt/gt^* mice injected with vehicle or psilocybin during a 10 min open field assay. n= WT Veh: 9, WT Psi: 9, *Scn2a^gt/gt^* Veh: 8, *Scn2a^gt/gt^* Psi: 8. Two-way ANOVA with Bonferroni multiple comparisons: ns, p>0.05. (D_i_) Time WT or *Scn2a^gt/gt^* mice injected with vehicle or psilocybin spent exploring the center of the open field chamber during a 10 min open field assay. n= WT Veh: 9, WT Psi: 9, *Scn2a^gt/gt^* Veh: 8, *Scn2a^gt/gt^* Psi: 8. Two-way ANOVA with Bonferroni multiple comparisons: ns, p>0.05 and **, p<0.01. (D_ii_) Number of times WT or *Scn2a^gt/gt^* mice injected with vehicle or psilocybin entered the center of the open field chamber during a 10 min open field assay. n= WT Veh: 9, WT Psi: 9, *Scn2a^gt/gt^* Veh: 8, *Scn2a^gt/gt^* Psi: 8. Two-way ANOVA with Bonferroni multiple comparisons: ns, p>0.05.

### *Scn2a*-deficient mice display a blunted head twitch response (HTR) to psilocybin compared to WT mice

To further evaluate the differential behavioral responses to Psi between WT and *Scn2a^gt/gt^* mice, we examined the HTR, a well-established behavioral hallmark of psychedelic action [40, 41]. This response, characterized by rapid side-to-side head movements, is primarily mediated by the 5-HT2A receptor [42, 43], and the potency of hallucinogens in the mouse HTR assay has been shown to correlate with their hallucinogenic potency in humans [44]. To evaluate Psi responsiveness, we injected Veh or Psi and counted the number of head twitches observed over 30 minutes (**Figure 3A**). We found that *Scn2a^gt/gt^* mice injected with Psi had fewer head twitches compared to the WT mice (WT: Veh: 5.125 ± 0.88, Psi: 24.25 ± 1.96, *Scn2a^gt/gt^*: Veh: 6.38 ± 0.83, Psi: 17.00 ± 2.56) (**Figure 3B**) and a decreased response to Psi (WT: 473.2 ± 38.31 %, *Scn2a^gt/gt^*: 266.7 ± 40.12 %) (**Figure 3C**). These findings, along with our previous behavioral results, suggest that alterations in serotonergic signaling may contribute to the differential responses to Psi observed in *Scn2a^gt/gt^* mice.

**Figure 3:**
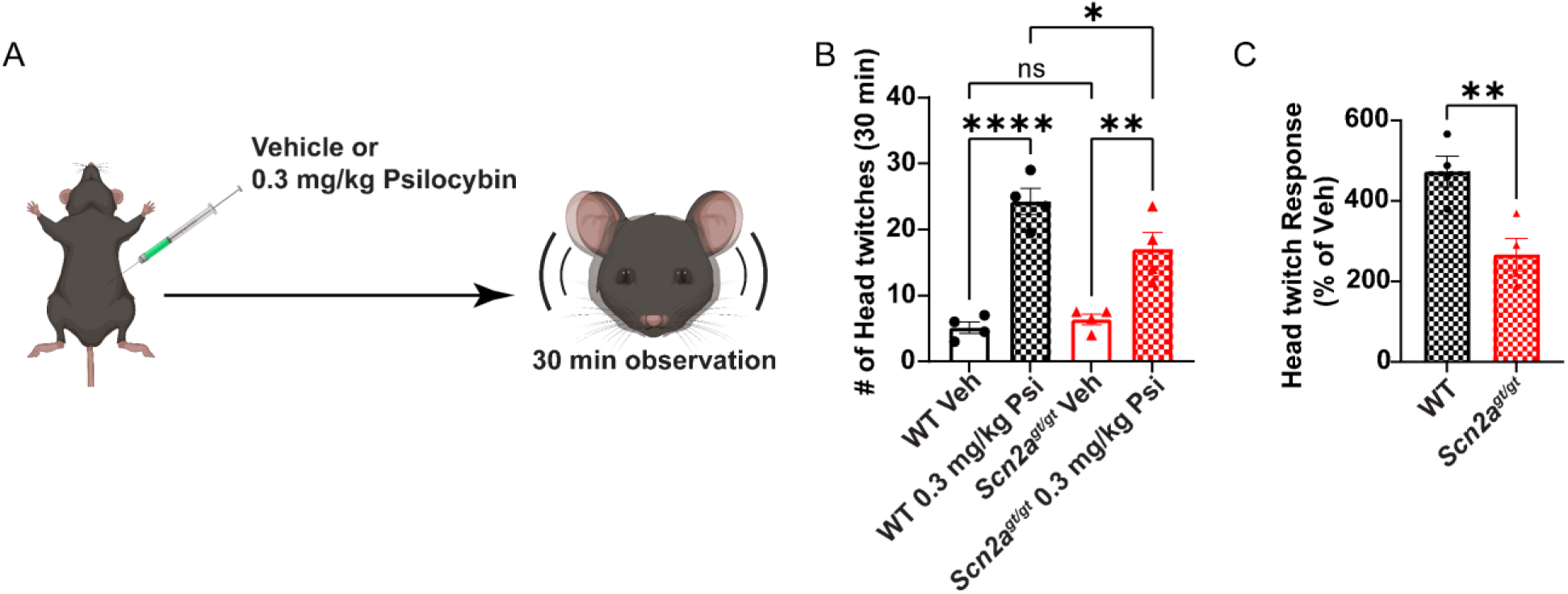
*Scn2a*-deficient mice display a blunted head twitch response to psilocybin compared to WT mice. (A) Schematic of experimental design. (B) Number of head twitches observed in WT or *Scn2a^gt/gt^* mice during the 30 minutes following vehicle or psilocybin administration. n= WT Veh: 4, WT Psi: 4, *Scn2a^gt/gt^* Veh: 4, *Scn2a^gt/gt^* Psi: 4. Two-way ANOVA with Bonferroni multiple comparisons: ns, p>0.05, *, p<0.05, **, p<0.01, and ****,p<0.0001. (C) Head twitch response to psilocybin in WT or *Scn2a^gt/gt^* mice, expressed as a percentage of the mean vehicle response for each genotype [(Number of head-twitches after psilocybin injection/average of head-twitches after vehicle injection for that genotype)*100]. n= WT Veh: 4, WT Psi: 4, *Scn2a^gt/gt^* Veh: 4, *Scn2a^gt/gt^* Psi: 4. Welch’s t-test: **, p<0.01.

### *Scn2a*-deficient mice exhibit altered serotonin receptor and transporter expression in the mPFC

To better understand the differential behavioral responses to Psi observed in *Scn2a^gt/gt^* mice, we next assessed the serotonergic system in the mPFC, a region implicated in both the head twitch response and social behavior [45, 46]. UHPLC-ECD analysis revealed no differences in 5-HT, 5-HIAA, or 5-HT turnover between *Scn2a^gt/gt^* and WT mice (5-HT: WT: 4.07 ± 0.21 ng/mg of protein, *Scn2a^gt/gt^*: 3.79 ± 0.16 ng/mg of protein, 5-HIAA: WT: 1.15 ± 0.10 ng/mg of protein, *Scn2a^gt/gt^*: 1.12 ± 0.08 ng/mg of protein, and 5-HT turnover: WT: 0.27 ± 0.010 [5-HIAA/5-HT], *Scn2a^gt/gt^*: 0.29 ± 0.01 [5-HIAA/5-HT]) (**Supplementary Figure 1A-C**). We next examined the expression of 5-HT receptors implicated in psychedelic signaling and social behavior [25, 26, 31, 33, 42, 43, 47–54]. qPCR analysis revealed that 5-HT1B mRNA expression was increased in *Scn2a^gt/gt^* mice (WT: 1.000 ± 0.1177, *Scn2a^gt/gt^*: 1.497 ± 0.08817), but the expression of 5-HT2A, 5-HT2C, 5-HT1A, 5-HT6, and 5-HT7 mRNA remained unchanged (**Supplementary Figure 2A-F**). These findings suggest that *Scn2a* deficiency is associated with relatively limited changes in serotonin receptor mRNA expression, despite the observed differences in behavioral responses to Psi.

Given that mRNA expression does not necessarily reflect protein abundance [55, 56], we next tested whether serotonergic alterations in *Scn2a^gt/gt^* mice occur at the protein level. Western blot analysis of 5-HT receptors in the mPFC revealed multiple changes in receptor expression, characterized by increased 5-HT1B (*Scn2a^gt/gt^*: 149.5 ± 16.6 % normalized to WT, p= 0.0299) (**Figure 4D**) and decreased 5-HT1A (*Scn2a^gt/gt^*: 79.79 ± 3.643 % normalized to WT, p= 0.0097), 5-HT2A (*Scn2a^gt/gt^*: 71.42 ± 6.256 % normalized to WT, p= 0.0411), 5-HT6 (*Scn2a^gt/gt^*: 43.06 ± 7.432 % normalized to WT, p= 0.0003), and 5-HT7 (*Scn2a^gt/gt^*: 67.71 ± 5.582 % normalized to WT, p= 0.0012) (**Figure 4C**, **4E**, **4G, and 4H**). However, 5-HT2C expression remained unchanged (*Scn2a^gt/gt^*: 95.46 ± 3.459 % normalized to WT, p= 0.5054) (**Figure 4F**). Given the widespread changes in 5-HT receptor expression, we also evaluated SERT and found that its expression was significantly decreased in *Scn2a^gt/gt^* mice (*Scn2a^gt/gt^*: 43.57 ± 10.31 % normalized to WT, p= 0.0015) (**Figure 4I)**. These findings identify alterations in serotonergic receptor and transporter abundance that may be associated with genotype-dependent behavioral responses to psilocybin.

**Figure 4:**
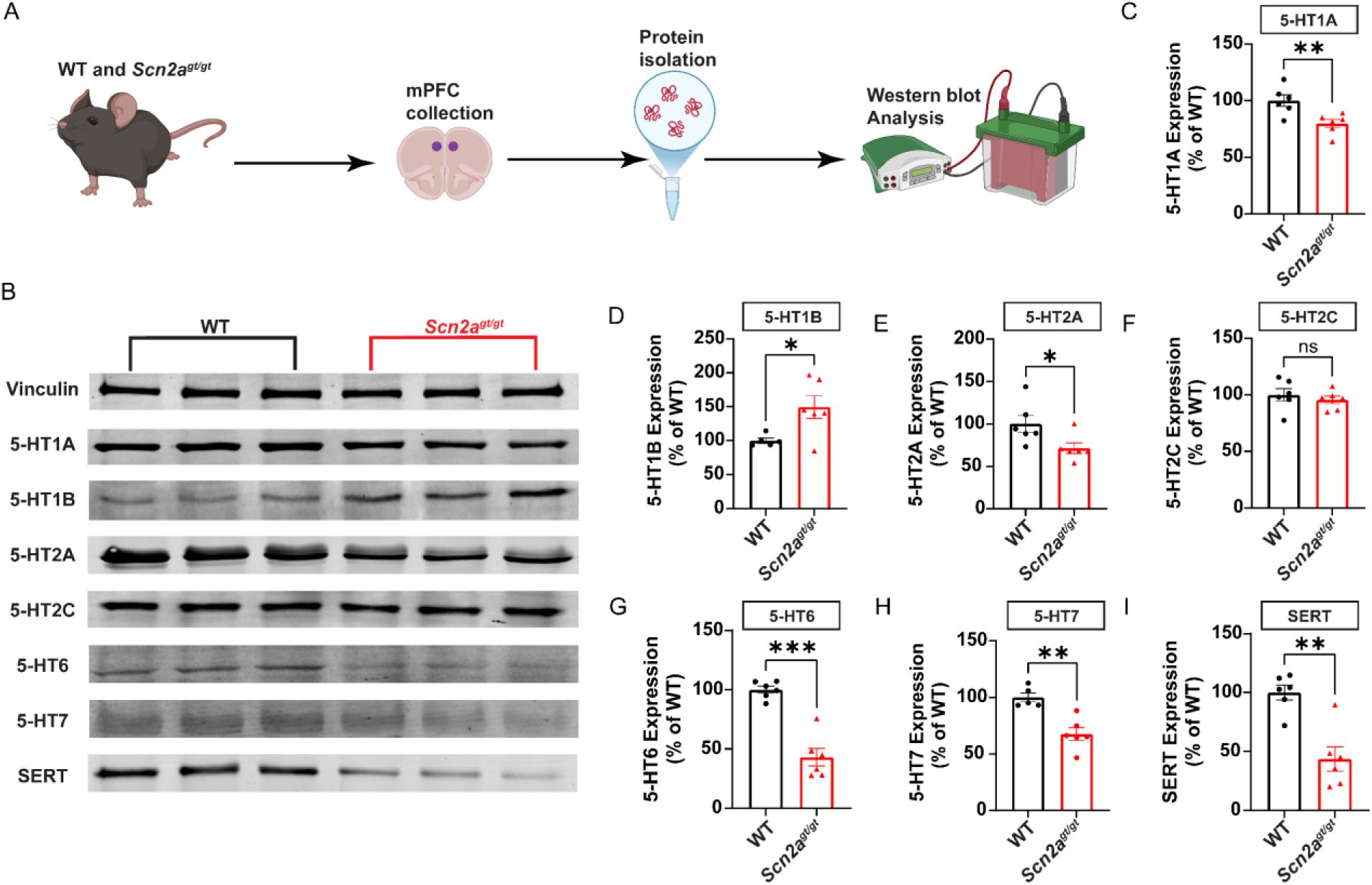
*Scn2a*-deficient mice exhibit altered serotonin receptor and transporter expression in the mPFC. (A) Schematic of experimental design. (B) Representative Western blot images of vinculin, 5-HT1A, 5-HT1B, 5-HT2A, 5-HT2C, 5-HT6, 5-HT7, and SERT from the mPFC of WT and *Scn2a^gt/gt^* mice. (C) Western blot analysis showing 5-HT1A expression in the mPFC of WT and *Scn2a^gt/gt^* mice. n= WT: 6, *Scn2a^gt/gt^*: 6. Welch’s t-test: **, p<0.01. (D) Western blot analysis showing 5-HT1B expression in the mPFC of WT and *Scn2a^gt/gt^* mice. n= WT: 5, *Scn2a^gt/gt^*: 6. Welch’s t-test: *, p<0.05. (E) Western blot analysis showing 5-HT2A expression in the mPFC of WT and *Scn2a^gt/gt^* mice. n= WT: 6, *Scn2a^gt/gt^*: 6. Welch’s t-test: *, p<0.05. (F) Western blot analysis showing 5-HT2C expression in the mPFC of WT and *Scn2a^gt/gt^* mice. n= WT: 6, *Scn2a^gt/gt^*: 6. Welch’s t-test: ns, p>0.05. (G) Western blot analysis showing 5-HT6 expression in the mPFC of WT and *Scn2a^gt/gt^* mice. n= WT: 6, *Scn2a^gt/gt^*: 6. Welch’s t-test: ***, p<0.001. (H) Western blot analysis showing 5-HT7 expression in the mPFC of WT and *Scn2a^gt/gt^* mice. n= WT: 5, *Scn2a^gt/gt^*: 6. Welch’s t-test: **, p<0.01. (I) Western blot analysis showing SERT expression in the mPFC of WT and *Scn2a^gt/gt^* mice. n= WT: 6, *Scn2a^gt/gt^*: 6. Welch’s t-test: **, p<0.01.

### *Scn2a*-deficient mice display elevated spontaneous firing of Dorsal Raphe Nucleus neurons

Given the observed alterations in serotonin receptor and SERT expression, we next examined whether neuronal activity in the DRN, a major source of serotonergic projections in the brain [57], was also affected by *Scn2a* deficiency. We found that DRN neurons in *Scn2a^gt/gt^* mice exhibited increased spontaneous firing compared with WT controls (WT: 1.97 ± 0.26 Hz, *Scn2a^gt/gt^*: 3.58 ± 0.34 Hz) (**Figure 5B-C**), accompanied by a leftward shift in the cumulative probability distribution of inter-event intervals (**Figure 5D**). These findings reveal altered DRN neuronal activity in *Scn2a*-deficient mice, suggesting that changes in serotonergic circuit activity may accompany the observed receptor alterations and potentially contribute to altered responses to psilocybin.

**Figure 5:**
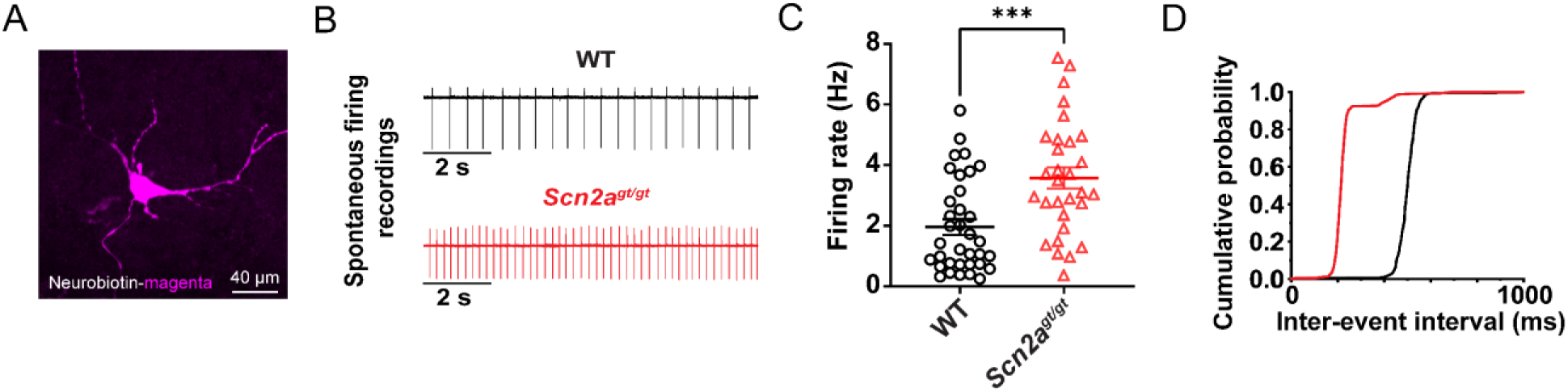
*Scn2a*-deficient mice display elevated spontaneous firing of Dorsal Raphe Nucleus neurons. (A) A representative Dorsal Raphe Nucleus neuron labeled with Neurobiotin. Scale bar, 40 µm. (B) Representative traces of spontaneous firing of DRN neurons in WT and *Scn2a^gt/gt^* mice. (C) Firing rate of DRN neurons in WT and *Scn2a^gt/gt^* mice. n= WT: 35 cells from 4 mice, *Scn2a^gt/gt^*: 30 cells from 4 mice. Unpaired t-test: ***, p<0.001. (D) Cumulative probability distribution of inter-event intervals for DRN neurons in WT and *Scn2a^gt/gt^* mice.

## Discussion

In this study, we demonstrate that Psi acutely increases social behavior in *Scn2a^gt/gt^* mice, an effect not observed in WT mice. Despite this behavioral improvement, *Scn2a^gt/gt^* mice exhibit a blunted HTR, suggesting altered serotonergic signaling. Interestingly, we found a discrepancy between 5-HT receptor mRNA and protein expression, revealing dysregulation of serotonergic receptors as a potential driver of altered Psi responsiveness. To evaluate a potential reason why 5-HT receptor changes in the mPFC are observed in *Scn2a^gt/gt^* mice, we evaluated the activity of DRN neurons. We found that *Scn2a^gt/gt^* mice exhibit increased spontaneous firing of DRN neurons. These findings suggest a broader multi-level disruption in serotonergic signaling, highlighting the importance of future research on the serotonergic system in *SCN2A*-related ASD.

Consistent with prior observations, *Scn2a^gt/gt^* mice exhibited social deficits at baseline [14]. Notably, an acute dose of Psi increased the sociability of *Scn2a^gt/gt^* mice to levels comparable to those of WT mice. This observation is consistent with published literature, where others have observed that psychedelics can increase social behavior in ASD mouse models [23, 58, 59]. More recently, a study demonstrated that Psi increased sociability in *Cntnap2* knockout mice but not WT mice, consistent with our observations [39]. Interestingly, we found that the prosocial effects of Psi were transient, and baseline behavior returned after 1 week. In contrast, Song et al. reported that the prosocial effects of Psi persisted for at least two weeks following administration [39]. One possible explanation for this difference is the reduced 5-HT2A expression observed in the mPFC of *Scn2a^gt/gt^* mice, a change that, to our knowledge, has not been reported in *Cntnap2* knockout mice. Previous studies have demonstrated that 5-HT2A receptors play an important role in Psi-induced structural plasticity [60, 61], suggesting that reduced receptor expression may influence the duration of Psi’s prosocial effects in *Scn2a^gt/gt^* mice.

Notably, the observation that psilocybin improved sociability despite an attenuated head twitch response suggests that these behavioral effects may involve distinct or partially overlapping serotonergic mechanisms. However, the contribution of 5-HT2A signaling to the prosocial response remains to be determined. The HTR is largely used as a proxy for 5-HT2A receptor activation following psychedelic administration. However, prior studies have demonstrated that additional 5-HT receptors can modulate both HTR and behavioral outcomes induced by psychedelics [26, 42, 43], indicating that psychedelic responses are shaped by the broader serotonergic system rather than a single receptor pathway. To evaluate if the behavioral changes observed in *Scn2a^gt/gt^* mice were due to altered serotonergic tone, we assessed baseline 5-HT and 5-HIAA levels in the mPFC. We found no significant difference between the levels of 5-HT, 5-HIAA, and 5-HT turnover in WT and *Scn2a^gt/gt^* mice. Previous studies investigating brain levels of 5-HT in ASD mouse models have shown inconsistent findings, suggesting that alterations in 5-HT availability are not standardized between ASD models [27].

Interestingly, we observed a discrepancy between 5-HT receptor mRNA and protein expression. Although this finding may initially appear counterintuitive, similar discrepancies have been reported in Alzheimer’s disease [55, 56]. This finding highlights the importance of interpreting mRNA data with caution, as mRNA abundance does not necessarily predict protein abundance [62]. Our Western blot analysis revealed that *Scn2a^gt/gt^* mice exhibit decreased expression of 5-HT1A, 5-HT2A, 5-HT6, and 5-HT7 in the mPFC. Our finding that *Scn2a^gt/gt^* mice have a reduced HTR is consistent with previous studies demonstrating the importance of 5-HT2A in this behavioral outcome following psychedelic administration [42, 63]. Reduced cortical 5-HT2A receptor binding has also been reported in individuals with ASD and associated with greater social communication difficulties in individuals with Asperger’s syndrome [64, 65]. Additionally, 5-HT1A agonism with 8-OH-DPAT has been shown to improve sociability in a valproic acid-induced rat model of ASD [51]. Beyond their involvement in behavioral regulation, serotonergic receptors also modulate neuronal activity. For example, 5-HT6 agonism has been shown to decrease excitatory synaptic transmission in prefrontal cortical pyramidal neurons [66]. Similarly, 5-HT7 receptors regulate neuronal excitability and seizure susceptibility [67], which may be particularly relevant given the high prevalence of seizures among individuals with ASD [68, 69]. We also observed increased 5-HT1B expression in the mPFC of *Scn2a^gt/gt^* mice, consistent with reports of increased cortical 5-HT1B binding in mice carrying an ASD-associated SERT variant [70]. Notably, 5-HT1B agonism has been shown to rescue social behavior across mouse models of ASD [35]. Together, our findings and previous studies highlight the importance of these serotonin receptors in regulating ASD-related behavior and neuronal excitability.

Our patch-clamp analysis revealed that *Scn2a^gt/gt^* mice have increased spontaneous firing of DRN neurons. This brain region is home to the largest population of serotonergic neurons that project to many different areas of the brain, including the mPFC [71]. Interestingly, previous studies have demonstrated that optogenetic activation of the DRN in *Shank3*^-/-^ mice results in prosocial effects [72], whereas inhibition of serotonergic projections from the DRN to the anterior cingulate cortex reduces social behavior [73]. However, despite having increased DRN neuronal activity, *Scn2a^gt/gt^* mice exhibit reduced sociability. One hypothesis is that *Scn2a^gt/gt^* mice exhibit hyperexcitability in DRN neurons to compensate for lower 5-HT release. In a prior study, we observed that *Scn2a^gt/gt^* mice exhibit decreased spontaneous and miniature EPSCs and IPSCs, suggesting reduced synaptic transmission [14]. Although UHPLC-ECD analysis revealed that the total tissue content of 5-HT in the mPFC is not significantly different from that of WT mice, this method does not capture changes in the dynamics of serotonin release in the mPFC.

Together, our findings reveal serotonergic dysfunction in *Scn2a^gt/gt^* mice and genotype-dependent behavioral responses to Psi. To our knowledge, this is the first study to characterize alterations in serotonergic receptor expression and DRN neuronal activity in a mouse model of *SCN2A*-related ASD. Furthermore, our findings demonstrate that psilocybin can acutely improve social deficits associated with *SCN2A* deficiency. The potential relevance of these findings extends beyond preclinical models, as a recent preprint reported differences in the effects of Psi on resting-state functional connectivity between autistic and non-autistic adults [74]. Although further research is needed to understand the mechanisms underlying these differences, our findings emphasize the importance of investigating serotonergic interventions in the context of *SCN2A*-related ASD.

## METHODS

### Experimental animals

All mice for experiments (WT (*Scn2a^+/+^*) and *Scn2a^gt/gt^*) were offspring of heterozygous (*Scn2a^+/gt^*) breeding pairs. Mice were group-housed with same-sex littermates (2-5 per cage). Mice were housed on a 12-h/12-h light/dark cycle with ad libitum access to food (2018S Teklad from Envigo) and reverse osmosis water. Cages were stored on vented racks and were filled with 1/8” Bed-o-cobb bedding (Anderson, Maumee, OH, USA). A >8 g shredded puck was provided as nesting material and enrichment. Adult mice (2-4 months) of both sexes were used for all behavioral and biochemical experiments. All experiments were approved by the Institutional Animal Care and Use Committee (IACUC).

### Genotyping

Mice were ear-punched at the time of weaning (P21-P28), and tissue samples were collected for genotyping. DNA was extracted by dissolving samples in 50 mM NaOH for 30 minutes at 95°C. Genotyping for the tm1a trapping cassette was then performed on the extracted DNA using polymerase chain reaction (PCR) with the following primers (forward 5′ to 3′: GAGGCAAAGAATCTGTACTGTGGGG, and reverse 5′ to 3′: GACGCCTGTGAATAAAACCAAGGAA). The PCR products were 240 bp (WT) and 340 bp (gt).

### Drug preparation and administration

Drugs were administered intraperitoneally. Psilocybin (0.3 mg/kg) (Cayman Chemical, 14041) was dissolved in saline. For the head twitch response, mice were injected immediately before testing. For open field and social assays, mice were injected 30 minutes prior to testing.

### RT-qPCR

Total mRNA from the mPFC was extracted using TRIzol reagent (Thermo Fisher Scientific, 15596018). RNA was reverse transcribed using a Maxima First Strand cDNA Synthesis Kit (Thermo Fisher Scientific, K1672). cDNA was subjected to quantitative PCR using Toyobo Thunderbird SYBR qPCR Mix (Toyobo Research Reagents, QPS-201) with the following primers: 5-HT1A (F: GAC AGG CGG CAA CGA TACT, R: CCA AGG AGC CGA TGA GAT AGTT), 5-HT1B (F: CGC CGA CGG CTA CAT TTAC, R: AGC GAT TAC AAA GGC GTT GGA), 5-HT2A (F: TAA TGC AAT TAG GTG ACG ACT CG, R: GAG GCT TCG GAA GTG TTA GCA), 5-HT2C (F: TCA ATA CCT CCG ATG GTG GAC, R: GGT GGC ATT GTG CAG TTT CTT), 5-HT6 (F: GCT GTG CGT GGT CAT CGTA, R: CAT CAG GTC CGA CGT GAA GAG), 5-HT7 (F: ATC TCG GTG TGC TTT GTC AAG, R: GCG ATG AAG ACG TTG CAG AAG), and Gapdh (F: AGT CAA GGC GAA TGG GAA G, R: AAG CAG TTG GTG GTG CAG GAT G) in a C1000 Touch PCR thermal cycler (Bio-Rad). Gapdh mRNA levels were used as an endogenous control for normalization using the ΔCt method [75].

### Western blot

mPFC tissue samples from WT and *Scn2a^gt/gt^* mice were collected and flash-frozen until use. Brain tissue was then sonicated in RIPA lysis buffer (Thermo Fisher Scientific, 89901) supplemented with protease and phosphatase inhibitors (Thermo Fisher Scientific, A32953). Samples were then centrifuged at 16,000 g for 20 min at 4°C. Protein concentration in the supernatant was determined by the bicinchoninic acid assay (Thermo Fisher Scientific). Proteins in 1 × sample buffer [62.5 mM Tris-HCl (pH 6.8), 2% (w/v) SDS, 5% glycerol, 0.05% (w/v) bromophenol blue] were denatured at 95°C for 5 min. For each sample, 35 μg total protein lysates was loaded onto the 8% sodium dodecyl sulfate-polyacrylamide (SDS-PAGE) gels and transferred onto a PVDF membrane (Millipore, IPFL00010). Blots were blocked in 5% nonfat milk in Tris-buffered saline and Tween 20 (TBST) for 1 h at room temperature and probed with the primary antibody 5-HT1B (Novus/Bio-techne, NB100-56350, 1:500), 5-HT7 (Novus/Bio-techne, NB100-56352, 1:500), 5-HT2A (Novus/Bio-techne, NBP2-86929, 1:500), 5-HT1A (Novus/Bio-techne, NBP2-21590, 1:500), 5-HT2C (Santa Cruz, sc-17797, 1:500), 5-HT6 (Novus/Bio-techne, NBP1-46557, 1:500), SERT (Alomone, AMT-004, 1:500), and (Vinculin: ThermoFisher Scientific, PA5-29688, 1:1000) in antibody diluent (LI-COR, 927-65001) overnight at 4°C. After overnight incubation, the blots were washed three times in TBST for 10 min, followed by incubation with secondary antibody 800CW Goat Anti-Rabbit (LI-COR, 926-32211, 1:5000) or 800CW Goat Anti-Mouse (LI-COR, 926-32210, 1:5000) for 1 h at room temperature. Following three cycles of 10 min washes with TBST, the immunoreactive bands were scanned and captured by the Odyssey® CLx Imaging System (LI-COR Biosciences) and quantitatively analyzed by densitometry with ImageJ software (NIH). Each sample was normalized to its respective vinculin control.

### Ultra-high-performance liquid chromatography (UHPLC) for 5-HT and 5-HIAA quantification

Brains from WT and *Scn2a^gt/gt^* mice were rapidly removed, and the mPFC was dissected out. mPFC tissue samples were homogenized through sonication in 0.1 M perchloric acid and centrifuged at 16,400 x g for 35 min at 4°C. The supernatant was filtered using a 0.22 µm Spin-X filter tube by centrifugation at 3,000 x g for 15 min at 4°C. The protein pellet was resuspended with 0.1 M NaOH overnight at 4°C for protein quantification. Samples were stored at -80°C until quantification. Neurotransmitter and metabolite quantification was performed using UPLC (Antec Scientific ALEXYS Neurotransmitters system 180.0091W) with an electrochemical detector (DECADE Elite) and a standard curve using area under the curve. The mobile phase (100 mM phosphoric acid, 100 mM citric acid, 0.1 mM EDTA, 0.6 g/L 1-octanesulfonic acid, 5% v/v acetonitrile, 5% v/v Methanol, pH 3.0) was used to separate the analytes flowing at 50 µL/min through a Waters ACQUITY UPLC BEH C18 Column (130Å, 1.7 µm, 1 mm X 100 mm) and detector set at 800 mV. The resuspended protein pellet was quantified using bicinchoninic acid assay (Thermo Fisher Scientific). 5-HT and 5-HIAA concentrations were normalized to the protein content (ng analyte/mg protein). 5-HT turnover (Eq. 1) was calculated using the following equation:

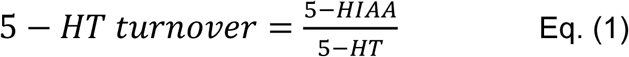

### Patch-clamp

#### Acute brain slice preparation

All chemicals were obtained from Sigma-Aldrich unless otherwise specified. Acute coronal brain slices containing the Dorsal Raphe Nucleus (DRN) were prepared from 3-8 month old WT and *Scn2a^gt/gt^* mice. Mice were deeply anesthetized with ketamine/xylazine (100/10 mg/kg, i.p., 0.1 mL/10 g body weight) before transcardial perfusion and decapitation. Brains were rapidly dissected into ice-cold slicing solution containing (in mM): 110 choline chloride, 2.5 KCl, 1.25 NaH_2_PO_4_, 25 NaHCO_3_, 0.5 CaCl₂, 7 MgCl₂, 25 glucose, 1 sodium ascorbate, and 3.1 sodium pyruvate (pH 7.4, 305–315 mOsm, bubbled with 95% O₂ and 5% CO₂).

Coronal slices (300 μm) containing the DRN were prepared using a vibratome (Leica VT1200S). Slices were incubated for 10 min at 33°C in slicing solution, then transferred to artificial cerebrospinal fluid (aCSF) containing (in mM): 125 NaCl, 2.5 KCl, 1.25 NaH_2_PO_4_, 25 NaHCO₃, 2 CaCl₂, 2 MgCl₂, and 10 glucose (pH 7.4, 305–315 mOsm, bubbled with 95% O_2_ and 5% CO_2_) for 10–20 min at 33°C before storage at room temperature for at least 30 min.

#### Spontaneous firing recordings

Slices were transferred to a recording chamber continuously perfused with aCSF at 32– 33°C (2–3 mL/min). DRN neurons were visualized using infrared differential interference contrast (IR-DIC) microscopy (Olympus BX-51WI) equipped with an IR-2000 camera (Dage–MTI).

Spontaneous neuronal firing was measured using the loose cell-attached patch-clamp technique to minimize perturbation of endogenous membrane properties. Glass pipettes (4–6 MΩ) were filled with an internal solution containing (in mM): 122 KMeSO_4_, 4 KCl, 2 MgCl_2_, 0.2 EGTA, 10 HEPES, 4 Na_2_ATP, 0.3 Tris-GTP, and 14 Tris-phosphocreatine (adjusted to pH 7.25 with KOH, 295–305 mOsm). Spontaneous action potentials were recorded for 3 min in gap-free mode without current injection.

Electrophysiological signals were acquired using an Axon MultiClamp 700B amplifier (Molecular Devices) and pClamp software 11.2, filtered at 2 kHz, and digitized at 30 kHz using an Axon Digidata 1550B plus HumSilencer digitizer (Molecular Devices).

Spontaneous firing frequency was calculated as the number of detected action potentials divided by recording duration. Inter-event intervals were calculated as the time between consecutive action potentials and plotted as cumulative probability distributions. Recordings were obtained from 35 neurons across 4 WT mice and 30 neurons across 4 *Scn2a^gt/gt^* mice.

#### Neuronal labeling

For morphological visualization, DRN neurons were labeled with Neurobiotin (0.1%– 0.2%) included in the pipette solution. Following recording, slices were fixed in 4% paraformaldehyde, washed with PBS, and incubated overnight at 4°C with fluorescently conjugated streptavidin.

### Open field

On the day of the test, mice were first acclimated to the test room for 20 minutes. Mice were then injected with saline or psilocybin. After 30 minutes, mice were placed in a square arena (40 cm x 40 cm x 40 cm, Maze Engineers) for 10 min at 60 lux. Distance traveled, velocity, center duration, and center frequency were recorded by EthoVision XT (Noldus).

### Three-chamber assay

Social preference was assessed using a standard three-chamber choice test with an empty cylinder and a cylinder containing a same-sex novel mouse of similar age and size (40.5 cm x 60 cm x 22 cm, Maze Engineers). Mice were habituated to the three-chamber apparatus one day before the test. On the day of the test, mice were first acclimated to the test room for 20 minutes. Mice were then injected with saline or psilocybin. After 30 minutes, mice were recorded for five minutes using EthoVision XT (Noldus). This assay was repeated 3 times with one week between each condition (vehicle->psilocybin->washout). Videos were then hand-scored by a blinded, well-trained researcher for the number of interactions a test mouse had with the novel mouse.

### Stranger interaction

On the day of the test, mice were first acclimated to the test room for 20 minutes. Mice were then injected with saline or psilocybin. After 30 minutes, mice were placed in a clean cage with a novel WT mouse of the same sex, age, and size. Social behaviors (e.g., sniffing, following, rearing towards the novel mouse) were scored manually by a blinded researcher over a 10-minute session. This assay was repeated 3 times with one week between each condition (vehicle->psilocybin->washout).

### Head twitch response

Mice were injected with vehicle or psilocybin and recorded for 30 minutes. The number of head twitches was counted by two blinded, well-trained researchers who reviewed recordings at one-quarter of the original speed. The counts from both researchers were averaged for each mouse.

### STATISTICAL ANALYSES

Normality and equality of variance were assessed for all data using GraphPad Prism. Outliers were defined a priori as values more than two standard deviations from the mean of their respective experimental group. All data were analyzed blinded to genotype. For comparisons between two groups, an unpaired t-test or Welch’s t-test (parametric) was used. For comparisons involving two factors, two-way ANOVA with Bonferroni’s multiple comparisons test was used. For repeated measurements involving two factors, two-way repeated-measures ANOVA with Bonferroni’s multiple comparisons test was used.

Post hoc comparisons were carried out only when the primary measure showed statistical significance. All data were expressed as mean ± SEM, with statistical significance determined at p-values <0.05. In all figures, p>0.05 is represented as not significant (ns), p<0.05 is represented as (*), p<0.01 is represented as (**), p<0.001 is represented as (***), and p<0.0001 is represented as (****).

## ACKNOWLEDGMENTS

The research reported in this publication was supported by the NINDS of the NIH (R01NS117585, R01NS123154, R56NS144233 to Y.Y.). The authors gratefully acknowledge support from the Familie*SCN2A* Foundation. Brody Deming was supported by an NIH research supplement to promote diversity in health-related research (R01NS117585-S1). Behavioral assays were performed using the Purdue Animal Behavior Core (Dr. Wendy Koss).

## AUTHOR CONTRIBUTIONS

B.D. designed research; B.D., J.Z., J.B.L., and K.F. performed experiments; B.D., J.Z., M.H., K.F., and D.R. analyzed data; A.A. and R.R. assisted with project administration; B.D. and Y.Y. wrote the manuscript with input from J.C.

## RESOURCE AVAILABILITY

Further information and requests for resources and reagents should be directed to and will be fulfilled by the lead contact, Yang Yang.

## Supplementary Information

**Supplementary Figure 1:**
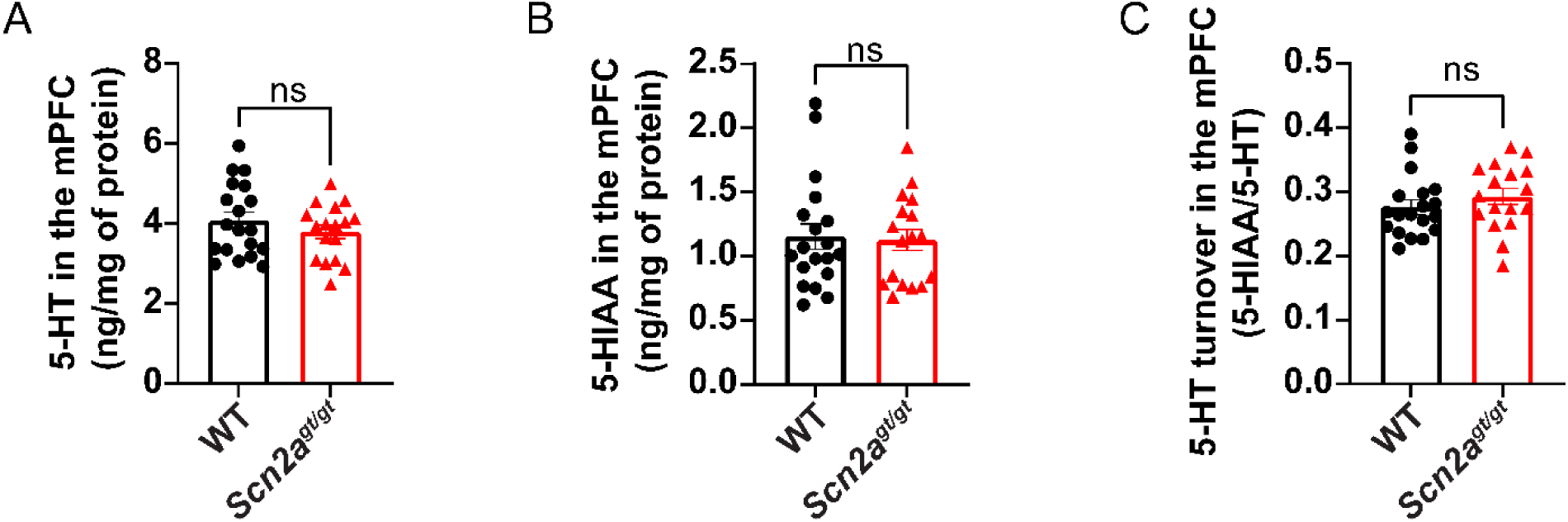
5-HT and 5-HIAA levels in the mPFC of WT and *Scn2a*-deficient mice. (A) Total mPFC concentration of 5-HT in WT and *Scn2a^gt/gt^* mice. n= WT: 19, *Scn2a^gt/gt^*: 17. Welch’s t-test: ns, p>0.05. (B) Total mPFC concentration of 5-HIAA in WT and *Scn2a^gt/gt^* mice. n= WT: 19, *Scn2a^gt/gt^*: 17. Welch’s t-test: ns, p>0.05. (C) mPFC 5-HT turnover (5-HIAA/5-HT) of WT and *Scn2a^gt/gt^* mice. n= WT: 19, *Scn2a^gt/gt^*: 17. Welch’s t-test: ns, p>0.05.

**Supplementary Figure 2:**
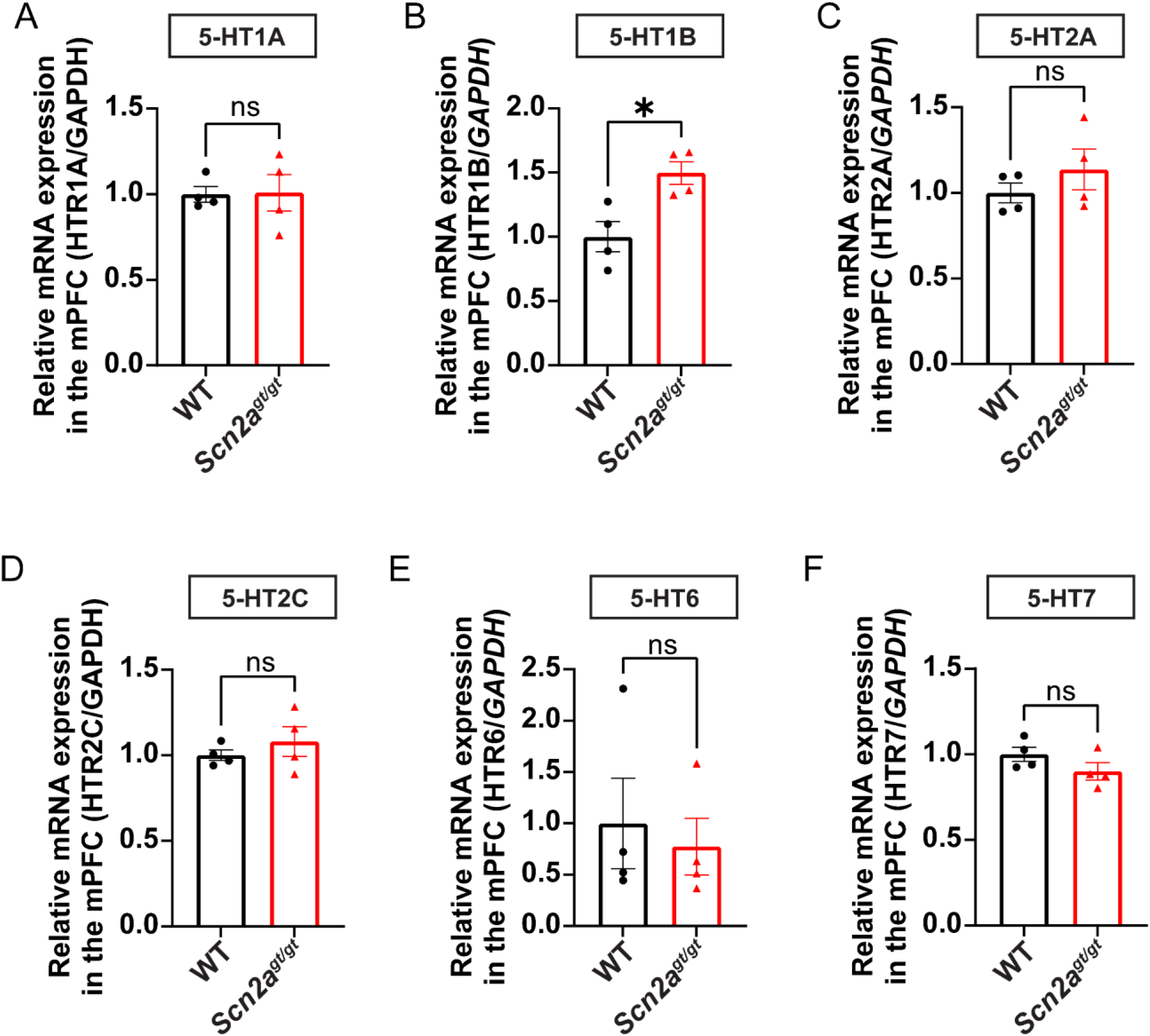
Serotonin receptor mRNA expression in the mPFC of WT and *Scn2a*-deficient mice. (A) Quantification of relative mRNA expression of 5-HT1A in the mPFC of WT and *Scn2a^gt/gt^* mice. n= WT: 4, *Scn2a^gt/gt^*: 4. Welch’s t-test: ns, p>0.05. (B) Quantification of relative mRNA expression of 5-HT1B in the mPFC of WT and *Scn2a^gt/gt^* mice. n= WT: 4, *Scn2a^gt/gt^*: 4. Welch’s t-test: *, p<0.05. (C) Quantification of relative mRNA expression of 5-HT2A in the mPFC of WT and *Scn2a^gt/gt^* mice. n= WT: 4, *Scn2a^gt/gt^*: 4. Welch’s t-test: ns, p>0.05. (D) Quantification of relative mRNA expression of 5-HT2C in the mPFC of WT and *Scn2a^gt/gt^* mice. n= WT: 4, *Scn2a^gt/gt^*: 4. Welch’s t-test: ns, p>0.05. (E) Quantification of relative mRNA expression of 5-HT6 in the mPFC of WT and *Scn2a^gt/gt^* mice. n= WT: 4, *Scn2a^gt/gt^*: 4. Welch’s t-test: ns, p>0.05. (F) Quantification of relative mRNA expression of 5-HT7 in the mPFC of WT and *Scn2a^gt/gt^* mice. n= WT: 4, *Scn2a^gt/gt^*: 4. Welch’s t-test: ns, p>0.05.

